# DNA METHYLTRANSFERASE 3 (MET3) is regulated by Polycomb Group complex during Arabidopsis endosperm development

**DOI:** 10.1101/2021.06.28.450243

**Authors:** Louis Tirot, Pauline E. Jullien

## Abstract

Complex epigenetic changes occur during plant reproduction. These regulations ensure the proper transmission of epigenetic information as well as allowing for zygotic totipotency. In *Arabidopsis*, the main DNA methyltransferase is called MET1 and is responsible for methylating cytosine in the CG context. The *Arabidopsis* genome encodes for three additional reproduction-specific homologs of *MET1*, namely *MET2a, MET2b* and *MET3*. In this paper, we show that the DNA methyltransferase *MET3* is expressed in the seed endosperm and its expression is further restricted to the chalazal endosperm. *MET3* is biallelically expressed in the endosperm but displays a paternal expression bias. We found that *MET3* expression is regulated by the Polycomb complex proteins FIE and MSI1. Seed development is not impaired in *met3* mutant, and we could not observe significant transcriptional changes in *met3* mutant. Interestingly, we found that *MET3* regulates gene expression in a Polycomb mutant background suggesting a further complexification of the interplay between H3K27me3 and DNA methylation in the seed endosperm.

**Key message:** The DNA METHYLTRANSFERASE MET3 is controlled by Polycomb group complex during endosperm development.

## Introduction

Sexual reproduction in Angiosperm is initiated by a double fertilization event. Fertilization of the haploid egg cell by one of the sperm cells gives rise to the diploid embryo whereas fertilization of the homodiploid central cell gives rise to the triploid endosperm (Berger 2003; Costa et al. 2004). The endosperm represents a nourishing tissue supporting embryo growth and is therefore key for proper seed development. Cell divisions in the endosperm are initiated very rapidly following fertilization. These divisions are initially occurring without cellularization and form a syncytium that will later cellularize (Brown et al. 1999, 2003; Boisnard-Lorig et al. 2001). An additional peculiarity of the endosperm, beyond its triploid syncytial nature, is being the seat of interesting epigenetic phenomena and complex epigenetic regulation.

Endosperm development and cellularization are indeed regulated by the FIS Polycomb group complex known to mediate Histone H3 Lysine 27 tri-methylation, a key silencing epigenetic mark. Some of the key members of the FIS Polycomb group complex (FIS-PcG) are MEA, FIS2, FIE, and MSI1 (Chaudhury et al. 1997; Luo et al. 1999; Kiyosue et al. 1999; Yadegari et al. 2000; Köhler et al. 2003; Guitton et al. 2004). In mutants affecting those genes, the endosperm fails to cellularize, resulting in an arrest of embryo development and eventually seed abortion. Additionally, several genes were found to be imprinted, *i.e*. only one parental allele is expressed whereas the other is epigenetically silent, in the endosperm. This is the case, for example, of genes such as *FIS2, FWA, MEA* or *PHE1* (Kinoshita et al. 1999, 2004; Luo et al. 2000; Köhler et al. 2005; Jullien et al. 2006). The silencing of those endosperm imprinted genes relies principally on two epigenetic mechanisms: either regulation by the FIS-PcG itself like for *MEA* or *PHE1* or silencing by DNA methylation like for *FWA* and *FIS2* (Jullien and Berger 2009; Gehring 2013; Batista and Köhler 2020). Another epigenetic singularity of the endosperm is of being relatively hypomethylated compared to other plant tissues (Gehring et al. 2009; Hsieh et al. 2009). This hypomethylation is due in part to the activity of a DNA glycosylase called DEMETER (Choi et al. 2002; Gehring et al. 2006; Hsieh et al. 2009) and likely also to the low expression of canonical actors of the DNA methylation pathway (Jullien et al. 2012).

DNA methylation is a key epigenetic mark regulating gene expression and protecting genome integrity by repressing transposons. In plant genomes, DNA methylation is found in three cytosine contexts: CG, CHG and CHH (where H is any base except C). Methylation on these different contexts relies on specific DNA methyltransferases. DNA methylation on CG sites relies on maintenance DNA METHYLTRANSFERASE (MET) where the main ubiquitous enzyme is MET1. DNA methylation on CHG sites relies on CHROMOMETHYLASE3 (CMT3) and an interplay with histone methylation (Lindroth et al. 2001). DNA methylation on CHH site, due to its non-symmetrical nature, relies on the constant *de novo* methylation pathway involving small RNA molecules as well as DOMAIN REARANGED METHYLTRANSFERASE2 (DRM2) (Cao and Jacobsen 2002). In centromeric sequences, CHH methylation also relies on CHROMOMETHYLASE2 (CMT2) (Stroud et al. 2013).

Although we know a lot about the main actors of this pathway, the *Arabidopsis’s* genome encodes multiple copies of DNA methyltransferase genes (4 *METs*, 3 *CMTs* and 3 *DRMs*) some of which might have a more complex or similar function in discreet cell types. For example, *CMT1* (the third CHROMOMETHYLASE encoded by the *Arabidopsis* genome) is principally detected in reproductive tissue (Henikoff and Comai 1998; Klepikova et al. 2016) and the reconstitution of a full-length transcript relies on the splicing out of a transposable element situated in its 13^th^ exon (Yadav et al. 2018). The *DOMAIN REARANGED METHYLTRANSFERASE1* (*DRM1*) seems to also solely play a role in reproductive tissue, where a redundancy between *DRM1* and *DRM2* was observed in the early embryo (Jullien et al. 2012). Similarly, data concerning the potential function of non-canonical *METs* are scarce. *MET2a* and *MET2b* are detected in the central cell, but their function is unknown (Jullien et al. 2012). Nonetheless, correlative evidence suggest *MET2a* might be important to regulate transposon reactivation in wild *Arabidopsis* accessions (Quadrana et al. 2016) and involved in fungal response (Salvador-Guirao et al. 2018).

As mentioned, little is known about the DNA methyltransferase *MET3. MET3* is also named *MATERNAL EFFECT EMBRYO ARREST 57* (*MEE57*) as a transposon insertion associated with the *MET3* locus led to an arrest in endosperm development (Pagnussat et al. 2005). *MET3* is also reported to be the sole *MET* expressed in the endosperm (Jullien et al. 2012). Here, we show that *MET3* is specifically expressed in the endosperm in a biallelic fashion with a paternal bias. *MET3* expression is controlled by the FIS-PcG complex. Despite the initial report of a seed arrest phenotype in the *mee57* line, we did not see any seed developmental phenotype in two independent *met3* mutant alleles. Additionally, we could not see major changes in the seed transcriptome of *met3* mutant. Nevertheless, we could see an effect on the seed transcriptome in a *fie* mutant background suggesting that *MET3* might interplay with PcG gene regulation in the developing endosperm.

## Material and Methods

### Plant Materials, Growth Conditions and Genotyping

The wild-types Col-0 and Gr-1, the *MET3* mutant lines *met3-3* (GABI404F04), *met3-4* (GABI659H03) and PRC2 mutant lines *fie-362* (GABI_362D08) (Bouyer et al. 2011) and *msi1* (SAIL_429_B08) were provided by the Nottingham Arabidopsis Stock Center (NASC). *pMET3:H2B-RFP* line was previously described (Jullien et al. 2012). After sowing, plants were stratified in the dark at 4°C between 2 and 4 days. Plants were germinated and grown in growing chambers under long-day conditions (16h light 22°C / 8h dark 18°C). For the transmission analysis (Fig. S5a), plants were grown on Murashige and Skoog (MS1/2) media agarose plate in long-day conditions for 12 days before genotyping. Primers for genotyping are listed in Table S1.

### Microscopy and phenotype observation

DIC seed phenotype and GUS observations were done using a Leica DM2000 as previously described (Jullien et al. 2006). For seed development observation and counting (Fig. 4a), plants were synchronized by removing all open flowers from the inflorescences. After 6 days, the two first siliques situated above the previously removed flowers were picked for each inflorescence. We refer to this stage as 6 days after synchronization (6 DAS). Siliques were dissected, cleared using chloride hydrate solution and mounted on a slide for observation. *pMET3:H2B-tdTomato* and *pMET3:H2B-RFP* reporter lines were imaged using a laser scanning confocal microscope (Leica SP5). When necessary, brightness and contrast were uniformly modified by using ImageJ.

### Cloning and transformation

*pMET3:H2B-tdTomato* and *pMET3:H2B-GUS* were generated using the Gateway Cloning System (Invitrogen). All PCR fragments were amplified by PCR using the Phusion High-Fidelity DNA Polymerase (Thermo). Primer sequences used for cloning can be found in Table S1. All plasmids were transformed into wild-type Columbia-0 plants the by floral dipping method (Clough and Bent 1998). At least ten transgenic lines were analyzed per construct, which showed a consistent fluorescence expression pattern. An Illustration of the constructs can be found in Fig. S2a.

### RNA extraction, qPCR & RT-PCR

Total RNAs were extracted using RNeasy Plant Minikit (Qiagen). All samples were treated with DNase I (ThermoScientific) at 37°C for 30 minutes. DNAse I was subsequently inactivated by the addition of EDTA and heat treatment (65°C for 10 minutes). First-strand cDNAs were synthesized using between 500 and 1000 ng of DNase treated total RNAs as a template. The RT reaction was performed using either Maxima First Strand cDNA Synthesis Kit (Fig. 1a, Fig. 2a) (ThermoScientific), containing both oligo-dT and random hexamer primers, or RevertAid RT Reverse Transcription Kit (Fig. 3e, Fig. S4b) (ThermoScientific), containing only oligo-dT primers. The qPCR reactions were performed with a QuantStudio 5 thermocycler (ThermoScientific) using SYBR green (KAPA SYBR FAST qPCR Master Mix or ORA qPCR Green ROX H Mix). The qPCR mix was prepared according to the manufacturer’s protocol. An RNA equivalent of 25 ng of cDNA templates was used for each reaction. The qPCR program was as follow: 95 °C for 3 minutes followed by 45 cycles of 95 °C for 5 seconds and 60 °C for 30 seconds. *ACTIN2* (*AT3G18780*) expression was used to normalize the transcript level in each sample. For each condition, RNA abundance of target genes was calculated from the average of three independent biological replicates with three qPCR technical replicates. Real-time PCR primers used in this study are listed in Table S1. For the allele-specific RT-PCR (Fig.2a), cDNAs were amplified for 22 cycles for *ACT2* primers and 35 cycles for *MET3* specific primers. Half of the *MET3* PCR product was digested for 1h30 at 37°C with XbaI restriction enzyme (Sigma-Aldrich). *ACT2* amplification was used as a control.

**Figure 1.**
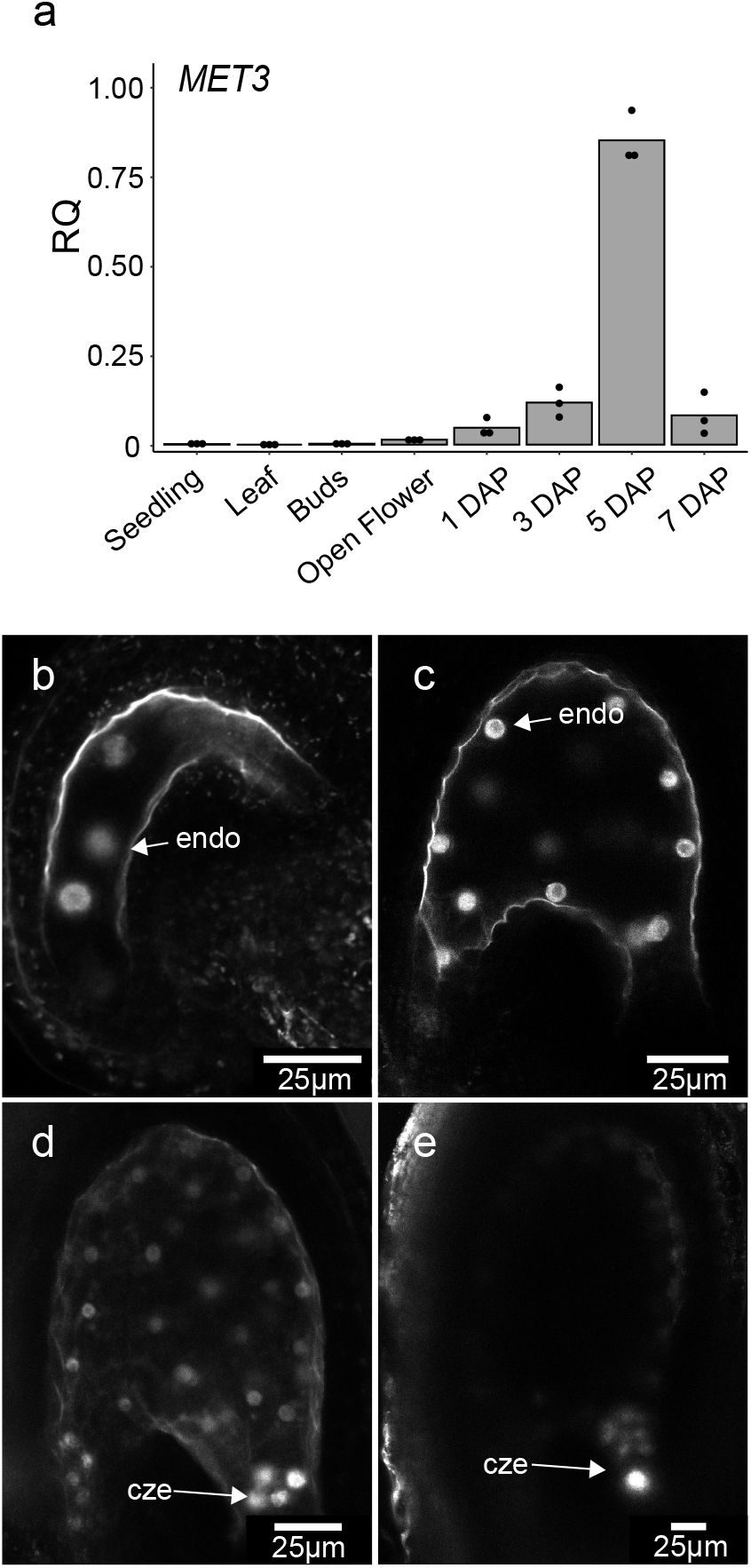
MET3 expression pattern. (a) *MET3* expression measured by RT-qPCR. DAP = Day After Pollination. The histogram displays the mean, and each dot represents a biological replicate. *ACT2* is used as a normalizer. RQ = Relative Quantification (b-e) Single-plan Confocal images representing the expression of *pMET3:H2B-tdTomato* construct in *Arabidopsis* 1 Day After Synchronization (DAS) seed (b), 2 DAS seed (c), 3 DAS (d) and 5 DAS (e). cze = chalazal endosperm, endo = endosperm nuclei. Scale bars represent 25μm.

**Figure 2.**
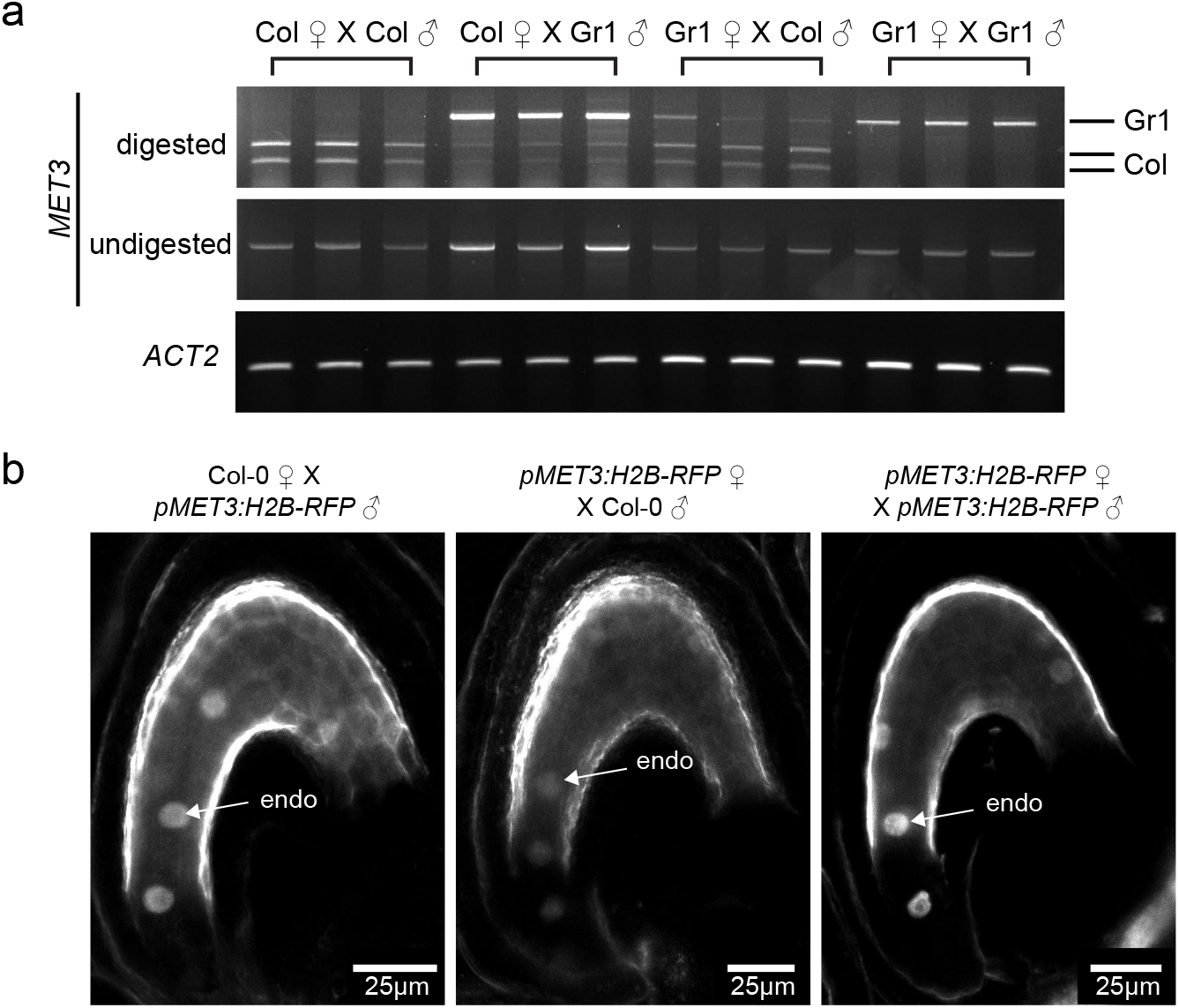
MET3 is biallelically expressed with a paternal bias. (a) Allele specific RT-PCR of *MET3* parental expression in 5DAP silique samples. The *XbaI* restriction enzyme digests the Col-0 *MET3* transcript but not the Gr-1 transcript. *ACT2* is used as loading control. (b) Single-plan confocal images of *pMET3:H2B-RFP* parental expression in the endosperm of 1 DAP seeds. endo = endosperm nuclei. Scale bars represent 25μm. ♂ symbol indicates the genotype of the male parent while ♀ indicates the genotype of the female parent.

### RNA sequencing and Bioinformatics

Total RNAs were extracted and DNAseI treated as previously mentioned. mRNA libraries were prepared and sequenced by Novogene (https://en.novogene.com/). Bioinformatic analyses were performed on the Galaxy web platform (https://usegalaxy.org) (Afgan et al. 2018). Our Galaxy workflow including the exact parameters and tool versions used can be downloaded on https://usegalaxy.org/u/pej/w/pejrnaseq and be freely reused. Briefly, Paired-end raw mRNA sequencing reads were controlled using FastQC (Galaxy version 0.72) and trimmed using Trimmomatic (Galaxy version 0.36.6) (Andrews 2010; Bolger et al. 2014). Clean reads were aligned to the *Arabidopsis thaliana* TAIR 10 genome assembly using HISAT2 (Galaxy version 2.1.0+galaxy4) (Kim et al. 2015). Aligned sequencing reads were assigned to genomic features using featureCounts (Galaxy version 1.6.2) (Liao et al. 2014). Differential expression was analyzed using DESeq2 default parameters (Galaxy Version 2.11.40.6+galaxy1) (Love et al. 2014). Differentially expressed genes (DEGs) were defined by an absolute logFC > 2 and an FDR < 0.05. Gene ontology (GO) enrichment analysis has been performed on PANTHER (Mi et al. 2019) and visualized using REVIGO (Supek et al. 2011). All plots have been generated using R-studio (www.rstudio.com). Raw data are deposited on the European Nucleotide Archive under reference PRJEB46544 (http://www.ebi.ac.uk/ena/data/view/PRJEB46544).

## Results

### MET3 is expressed biallelically with a paternal bias in the endosperm

Our previous analysis has shown that *MET3* is expressed in the endosperm of developing *Arabidopsis* seeds (Jullien et al. 2012). However, the detail and exclusivity of its expression pattern remain to be investigated. To get a better characterization of *MET3* expression pattern, we performed a qPCR of *MET3* transcript in different tissue types of wild-type Col-0 (Fig. 1a). Our result shows that *MET3* is principally expressed in siliques and its expression peaks at 5 Days After Pollination (DAP). These results could be confirmed using publicly available transcriptome datasets (Fig. S1a) (Klepikova et al. 2016). To gain a better spatial and temporal characterization of *MET3* expression, we generated two new transcriptional *MET3* reporter constructs, encompassing 2kb of the *MET3* promoter driving either H2B-tdTomato or H2B-GUS (Fig. S2a). The analysis of the *pMET3:H2B-GUS* in different plant tissues confirmed the specificity of *MET3* expression to the seed endosperm (Fig. S1b-d). To characterize in detail the temporal expression of *MET3* in the endosperm, we performed confocal microscopy on the *pMET3:H2B-tdTomato* lines at different stages of seed development. We could detect *pMET3:H2B-tdTomato* expression from as early as the four nuclei stage endosperm (Fig. 1b). *pMET3:H2B-tdTomato* remains express throughout the endosperm (Fig. 1c) until the globular stage of embryo development where its expression starts to be higher in the chalazal endosperm and chalazal cyst (Fig. 1d). At later stages, *pMET3:H2B-tdTomato* expression is limited to the chalazal endosperm and chalazal cyst (Fig. 1e). From 7 DAP, *pMET3:H2B-tdTomato* expression can no longer be detected. A similar expression pattern was observed with *pMET3:H2B-GUS* (Fig. S2b) and *pMET3:H2B-RFP* (Fig. S2c) as well as online transcriptomic data (Fig. S2d) (Belmonte et al. 2013). MET3 protein expression and localization could not be determined as we, so far, failed in expressing a fluorescently tagged MET3 protein in *Arabidopsis* (LT personal communication).

Such endosperm expression pattern is common in imprinted genes, like *FWA*, *FIS2*, *MEA* or *PHE1* (Kinoshita et al. 1999, 2004; Köhler et al. 2005; Jullien et al. 2006). In order to examine if *MET3* is biallelically or mono-allelically expressed, we performed allele-specific RT-PCR. We are making use of a Short Nucleotide Polymorphism (SNP) consisting of a substitution from a C to a T within *MET3* 9th exon in the Gr-1 ecotype which is abolishing a XbaI restriction site present in Col-0. We did reciprocal crosses using Col-0 and Gr-1 ecotypes and analyzed *MET3* parental expression at 5 DAP following XbaI digestion (Fig. 2a). We could observe, for both reciprocal crosses, bands corresponding to *MET3* transcript from Col-0 (505 and 329 bp) and from Gr-1 (834 bp) with a bias toward the paternal allele. This result shows that *MET3* is expressed from both maternal and paternal allele but displays a paternal bias of expression. *MET3* paternally biased expression was also observed using the *pMET3:H2B-RFP* transgene at 1 DAP (Fig.2b). Taking together our results shows that *MET3* expression is biallelic with a paternal bias and confined to the endosperm. *MET3* expression, initially throughout the endosperm, becomes restricted to the chalazal pole at later stages.

### MET3 expression is regulated by Polycomb group proteins

Beyond imprinted genes, *MET3* expression pattern is reminiscent of the expression of genes controlled by the endosperm FIS Polycomb group complex (PcG) composed of FIE, MSI1, FIS2 and MEA (Guitton et al. 2004). To investigate if *MET3* could be regulated by PcG, we introgressed a *MET3* transcriptional reporter into *fie* and *msi1* mutant background (Fig. 3a-d). We could observe increased *pMET3:H2B-RFP* reporter expression in 49% of the seeds (n=249) in *fie*/+ mutant and 44% (n=240) in *msi1*/+ mutant background characteristic of the maternal gametophytic effect of those mutations (Fig. 3d). In *msi1* and *fie* mutants, *pMET3:H2B-RFP* expression is higher and observed throughout the endosperm (Fig. 3b-c) whereas at the same developmental stage the expression of *pMET3:H2B-RFP* is already restricted to the chalazal pole in wild-type seeds (Fig. 3a). In order to confirm that the regulation of *MET3* by *MSI1* and *FIE* was not only restricted to our transgene, we performed a RT-qPCR of *MET3* expression in wild-type and mutant selected seeds at 10 DAP (Fig. 3e). We could observe a clear upregulation of *MET3* expression in both *fie* and *msi1* seeds compare to wild-type seeds (t-test p-value of 0.0543 and 0.0114 respectively). Using publicly available data, we could see that the upregulation of *MET3* is also observed in other PcG mutants (Fig. S3a-b). *MET3* upregulation was observed in silique samples of *clf* mutant (Fig. S3a) (Liu et al. 2016) and in *fis2* seeds (Fig. S3b) (Weinhofer et al. 2010).

**Figure 3.**
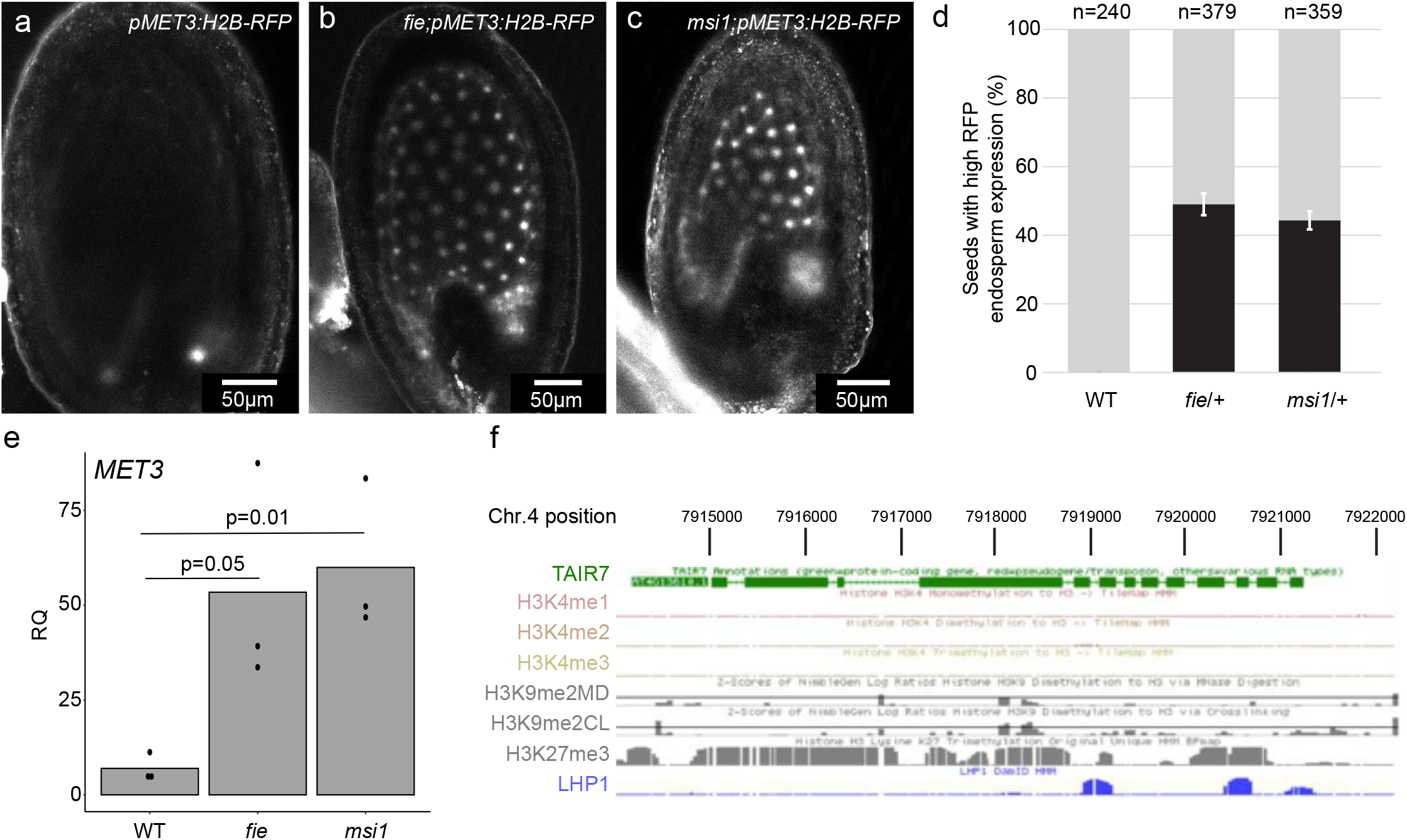
MET3 expression is controlled by FIE and MSI1 Polycomb proteins. (a-c) Confocal images representing the expression of *pMET3:H2B-RFP* construct in *Arabidopsis* wildtype (a), *fie* (b) and *msi1* (c) selfed seeds at 5 days after pollination (DAP). Scale bars represent 50 μm. (d) Proportion of seeds with high RFP signal in the endosperm at 5DAP in wild-type, *fie* and *msi1* mutants. Grey bars represent the seeds with a restricted RFP expression to the chalazal endosperm (as illustrated in (a)). Black bars represent the seeds with ectopic expression of *pMET3:H2B-RFP* throughout the endosperm (as illustrated in (b-c)). (e) *MET3* expression measured by RT-qPCR in 10 DAP selected seeds of wild-type, *fie* and *msi1*. The histogram displays the mean, and each dot represents a biological replicate. *ACT2* is used as a normalizer. (f) Snapshot showing that the *MET3* locus contains H3K27me3 and LHP1 but no H3K4me1, H3K4me2, H3K4me3 and H3K9me2 in seedlings. Data from http://epigenomics.mcdb.ucla.edu/H3K27m3/ (Zhang et al. 2007).

**Figure 4.**
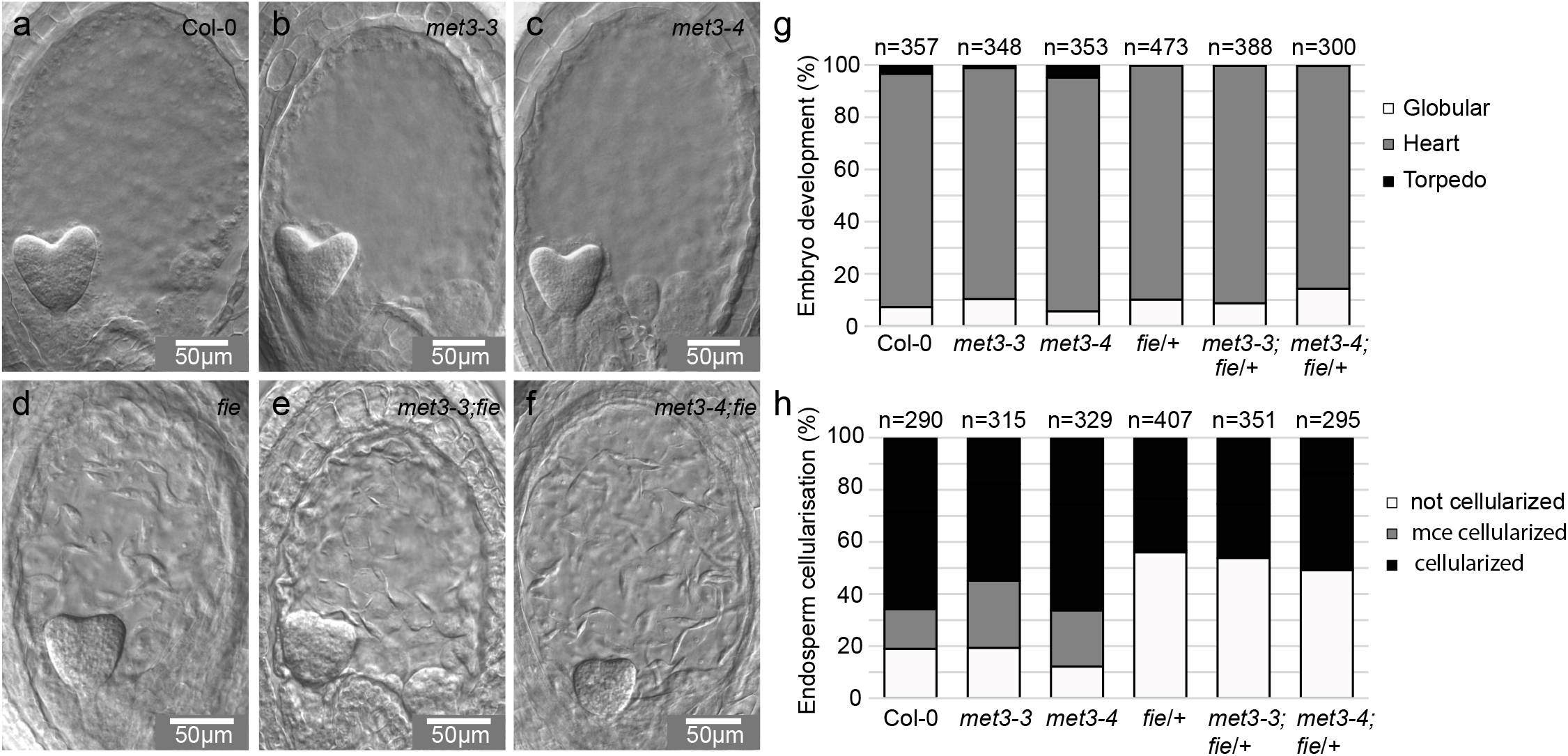
MET3 does not influence fie phenotype. (a-f) Seed developmental phenotype observed after clearing by Difference Interference Contrast (DIC) of wild-type (a), *met3-3* (b), *met3-4* (c), *fie*/+ (d), *met3-3;fie*/+ (e) and *met3-4;fie*/+ (f) at 6 days after synchronization (DAS). Scale bars represent 50 μm. (g-h) Quantification of the embryo developmental stages (g) and endosperm cellularization (h) in wild-type, *met3-3, met3-4, fie*/+, *met3-3;fie*/+ and *met3-4;fie*/+ at 6 DAS. mce = micropylar endosperm.

Subsequently, we wanted to know if the effect of the PcG complex was direct or indirect. PcG complexes are known to repress gene expression by tri-methylating the Lysine 27 of the Histone H3 tail (H3K27me3) inducing a closed chromatin state at the targeted loci. We, therefore, analyzed available H3K27me3 genome-wide Chromatin Immunoprecipitation (ChIP) data. We could see that the *MET3* locus is covered by H3K27me3 in *Arabidopsis* seedling samples (Fig. 3f) (Zhang et al. 2007). Additionally, using the RepMap2020 tool (Chèneby et al. 2020), we could observe H3K27me3 on the *MET3* locus in several independent ChIP experiments including some performed on endosperm tissue (Fig. S3c). We conclude that *MET3* expression is directly repressed by PcG complex induced H3K27me3 in the endosperm.

### MET3 does not affect seed development

Considering that *MET3* is specifically expressed in the developing endosperm during seed development, we then ask if *MET3* function influences seed and/or endosperm development. We characterized two mutant alleles from the GABI collection: *met3-3* (GABI_404F04) and *met3-4* (GABI_659H03) (Fig. S4a). The mutations are located on the 10^th^ and the 2^nd^ exon respectively and are expected to abolish *MET3* function. To confirm the downregulation of *MET3* in the mutants, we performed RT-qPCR. We could observe that *MET3* is downregulated in both mutant alleles (Fig. S4b). To investigate if the *met3* mutation could result in seed lethality we first investigated the presence of aborted seeds at the green seed stage (~12DAP). We could not see any significant seed abortion in both *met3-3* and *met3-4* alleles (Fig. S5b). To further confirm the absence of defects, we analyzed the transmission rate of the *met3* mutations in *met3-3*/+ and *met3-4*/+ selfed progeny. We could not observe any segregation distortion from the Mendelian ratio (Fig. S5a). Mutations affecting the main *Arabidopsis* DNA methyltransferase *MET1* display variation in seed size. We, therefore, investigated if *met3* mutants would display a seed size phenotype. We could not see any significant variation in seed size using both *met3* alleles (Fig. S5c-d). We conclude that *MET3* mutation does not severely impair seed development. Additionally, *met1* mutant are known to display increased developmental defect through generation (Mathieu et al. 2007). To assess potential transgenerational effect of the *met3* mutation we have maintained *met3* homozygotes mutants for five generations. However, we could not observe such increased developmental phenotype with *met3* mutants after five generation of inbreeding (*met3^G5^*) (Fig. S5e-g). The *met3^G5^* did not show difference when compared to wild-type either in term of rosette size (Fig. S5f) nor flowering time (Fig. S5g). We conclude that *MET3* mutations do not severely impair seed development and do not accumulate transgenerational developmental defects.

As shown above, *MET3* expression is regulated by PcG complex in the endosperm, we then analyzed if *MET3* mutation could influence the PcG *fie* mutant phenotype. To answer this question, we generated *fie*/+;*met3-3* and *fie*/+;*met3-4* double mutants and analyzed their seed development phenotype using DIC (Fig. 4a-f). In order to minimize the stress to the plant due to handling during emasculation and crossing, we used “synchronized seeds”. In practice, we remove the open flowers of the day, and we wait an x number of days before collecting two siliques above our cutting. This is allowing us to have age synchronized siliques without the physical disturbance of emasculation/pollination and is, therefore, closer to normal growth and fertilization. We are using the term Day After Synchronization (DAS). To our experience, 6 DAS is corresponding to around 4-5DAP. At 6 DAS, we could not see any delay in embryo development and endosperm cellularization comparing Col-0 to *met3-3* and *met3-4* mutants (Fig. 4a-c, 4g-h). In *fie*/+, we could clearly see the delayed endosperm cellularization characteristic of FIS-PcG mutants (Fig. 4d-f and h) (56% n=407) (Ohad et al. 1999; Sørensen et al. 2001). In the double mutants, we could not see variations in the quantification of the *fie*/+ phenotype for both embryo development (Fig. 4g) and endosperm cellularization (Fig. 4h). We conclude that *MET3* mutations do not influence *fie* mutant seed phenotype.

### Effect of MET3 mutation on the seed transcriptome

In order to investigate the potential effect of *met3* mutation on the seed transcriptome, we performed a mRNA deep sequencing experiment of 3 DAP (Fig. S6) and 10 DAS (Fig. 5a) seeds (*i.e*. seeds attached to the septum). We then compared the seed transcriptome of *met3* seeds to wild-type seeds. At 3 DAP, we could only detect one differentially expressed gene: *ESM1* (*AT3G14210*). *ESM1* was up-regulated in both *met3-3* and *met3-4* mutant seeds (Fig. S6a-b). At 10 DAS, we could also only see very minor changes to the seed transcriptome (Fig. 5a). We could identify only 16 differentially expressed genes with an absolute logFC>2 and FDR<0.05. These results show that reminiscent of the absence of seed phenotype in *met3* mutants, the seed transcriptome is also mostly unaffected by the *met3* mutation. As *MET3* expression is regulated by the FIS-PcG complex in the endosperm, we then ask if *MET3* mutation could influence the *fie* transcriptome. We, therefore, sequenced the transcriptome of 10 DAS seeds where we selected for *fie* mutant seeds under the dissecting microscope (white seeds). We compared the transcriptome of *fie* seeds to the transcriptome of *fie;met3-3* double mutant seeds. We could observe 87 differentially expressed genes (DEG) with an absolute logFC>2 and FDR<0.05 (Fig 5b). A Goterm enrichment analysis revealed that these genes are enriched for genes involved in pectin metabolism (GO:0045490, FDR=4.46E-10; GO:0045488, FDR=6.59E-09) and cell-wall related processes (GO:0042545, FDR= 8.97E-09; GO:0071555, FDR=4.87E-10, GO:0071554, FDR=5.67E-08) (Fig. 5c and Table S2). We then analyzed if among these 87 DEGs some are also modified in *fie* mutant compared to wild-type. We could find that a large proportion of either *met3* DEGs (Col vs *met3*, 11/16) and *met3;fie* DEGs (*fie* versus *met3fie*, 44/87) are miss-regulated in *fie* mutant seeds (Fig. 5d). We conclude that the *met3* mutation does not drastically change the seed transcriptome, but a set of genes are miss-regulated by both *met3 and fie mutants*.

**Figure 5.**
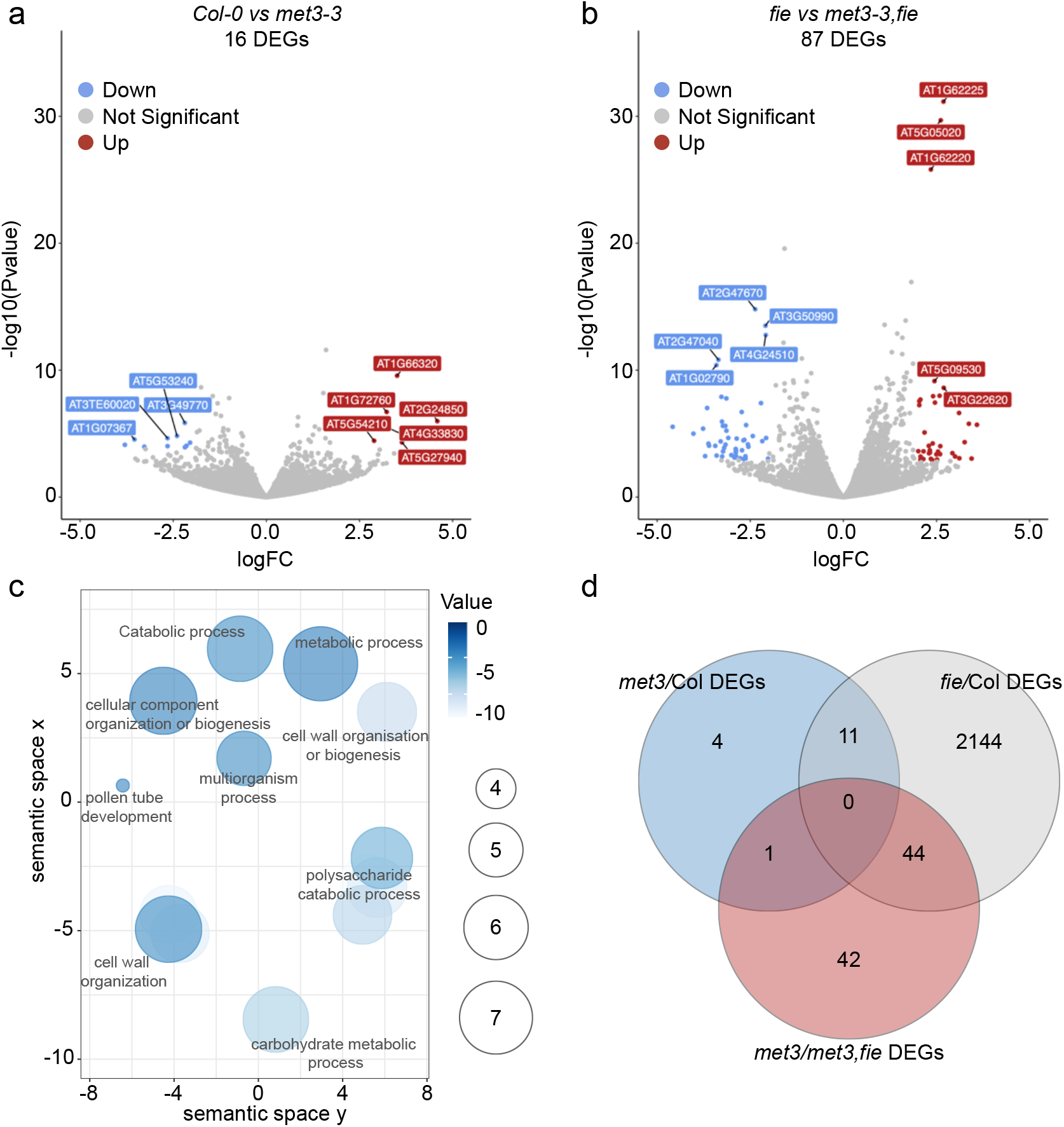
Transcriptome of met3 and met3;fie mutant seeds. (a-b) Volcano plot depicting differentially expressed genes (DEGs) comparing Col-0 to *met3-3* (a) and *fie* to *met3-3;fie* in 10 DAS seeds (b). Red dots represent up-regulated DEGs and blue dots represent down-regulated DEGs. The top 10 DEGs are annotated. We use a threshold of [absolute logFC > 2, FDR < 0.05]. (c) GO term analysis for the 87 DEGs obtained with the *fie vs met3-3;fie* contrast. (d) Venn diagram showing the overlapping DEGs between the different contrasts.

## Discussion

Our study highlights an additional connection between DNA Methylation pathways and Polycomb group H3K27 tri-methylation in the seed endosperm via the regulation of *MET3* by the FIS-PcG complex. *MET3* is specifically expressed in the endosperm, and its expression becomes restricted to the chalazal pole at later stages of endosperm development. In our study, using two independent insertion lines, *met3-3* and *met3-4*, we could not observe any major seed developmental phenotype. Both mutants show a significant decrease in *MET3* expression by qPCR suggesting they both represent knockout mutants. It was previously documented that a mutation affecting *MET3* named *mee57* displayed a strong seed developmental phenotype (Pagnussat et al. 2005). The *mee57* mutation shows an early maternal embryo and endosperm arrest. The discrepancy between our lines and the previously published *mee57* could have several causes: the mutagenesis method used (T-DNA versus transposition), the presence of additional genetic modifications, or a difference between the two ecotypes used, Columbia-0 in our case and Landsberg for *mee57*. If the latter is true, *MET3* function could vary between different *Arabidopsis* accessions.

In this work, we show that repression of *MET3* expression at later stages of endosperm development is linked to the direct action of the FIS-PcG complex on the *MET3* locus. *MET3* is over-expressed in PcG mutant seeds, such as *fie* and *msi1* mutant seeds. Interestingly, it was previously shown that *fie* mutant endosperm display higher CG methylation compared to wild-type endosperm and lower CHG and CHH (Ibarra et al. 2012). This increased CG methylation in *fie* mutant seems to be restricted to the endosperm as it is not observed in *fie* seedling or *fie* embryo methylome (Bouyer et al. 2017). We propose that higher expression of *MET3* in the *fie* endosperm could be the cause for the increase in CG methylation specifically in the endosperm. Indeed, in our study, we could not observe any change in *MET3* tissue expression pattern in PcG mutants but the increased expression is still restricted to the endosperm. In further studies, the Investigation of the endosperm methylome in *met3* and *met3;fie* mutants will allow to test this hypothesis. Additionally, further experiments are required to determine if MET3 is a functional DNA methyltransferase.

As we previously mentioned, *MET3* function could vary in between wild *Arabidopsis* accessions. Similarly, to CMT1, several SNPs can be found at the *MET3* locus in-between different ecotypes suggesting that *MET3* might not be fully functional in all of them. Additionally, looking closer at the *MET3* locus, we could detect the presence of a Class 2 DNA transposon (*AT4TE34810*) situated in the third intron of the *MET3* gene. In addition to PcG regulation, *MET3* expression and possibly function could be influenced by the presence or regulation of *AT4TE34810* like it is the case for *CMT1* transcript (Yadav et al. 2018). The study of *MET3* function and imprint in different accession could reveal more about its function in natural growth conditions. One hint of *MET3* potential function in Columbia-0 came from the study of methylome stability across several generations (Becker et al. 2011). In this study, methylome stability was investigated in several Col-0 selfed lineages (30 generations). The line accumulating the most methylation polymorphisms had concomitantly acquired a SNP in the *MET3* gene. It was therefore suggested that *MET3* could be the cause of such methylome instability (Becker et al. 2011; Schmitz and Ecker 2012). Analyzing the methylome of selfed *met3* mutants after 30 generations would allow to test this hypothesis. If true and taking into account the endosperm specificity of *MET3* expression, it would indicate that the endosperm methylome influences the embryonic methylome, a hypothesis that has stimulated a lot of interest over the last 10 years but remains to be demonstrated.

## Supporting information

supfig

## Acknowledgments

We would like to thank the following people for their help: Jasmine Sekulovski for support concerning plant growth, Diane Bonnet for technical support. We thank Mathieu Ingouff and Diane Bonnet for critical reading of the manuscript.

## Author contributions

PEJ conceived the study. LT performed the experiments. LT and PEJ analyzed the data. PEJ wrote the manuscript with the help of LT.

---

***Figure S1. MET3 is only expressed in seeds***

(a) Snapshot of the *MET3* expression pattern obtained from ebar using the data from Klepikova *et al*. (Winter et al. 2007; Klepikova et al. 2016)(b-d) The absence of *pMET3:H2B-GUS* expression observed in *Arabidopsis* seedling (b), stamen (c) and ovule (d).

***Figure S2. MET3 reporters and expression***

(a) Representation of *pMET3:H2B-GUS, pMET3:H2B-tdTomato* and *pMET3:H2B-RFP*. (b) *pMET3:H2B-GUS* expression observed in 1 DAP and 3 DAP seeds. (c) *pMET3:H2B-RFP* expression observed in 1 DAP and 3 DAP seeds. Scale bars represent 50 μm. (d) Snapshot of expression data from LCM dissected seeds from http://seedgenenetwork.net/plotprobe?name=254720_at (Belmonte et al. 2013).

***Figure S3. MET3 regulation by Polycomb group***

(a) *MET3* expression from an RNA-seq experiment in inflorescence, root, shoot and siliques from wild-type plants and *clf28* mutant plants, data from Liu *et al* 2016 (Liu et al. 2016). (b) *MET3* expression from transcriptomic data obtained from 3 DAP seeds using MicroArray comparing wild-type to *fis2-1* endosperm, data from Weinhofer *et al* 2010 (Weinhofer et al. 2010). (c) Snapshot of H3K27me3 ChIP data from several tissues including endosperm (colored in blue), flowers (colored in yellow) and various sporophytic (colored in green). Green line represent the presence of ChIP peaks of H3K27me3. Data from ReMap (Chèneby et al. 2020).

***Figure S4. Characterization of MET3 mutants***

(a) Representation of the *MET3* locus. Blue triangles are representing the T-DNA insertion site of both *met3-3* (GABI_404F04) and *met3-4* (GABI_659H03) (b) *MET3* expression measured by RT-qPCR in Col-0 and *met3* mutants seeds at 3DAP. The histogram displays the mean, and each dot represents a biological replicate. *ACT2* is used as a normalizer. 2 stars indicate a p value <0.01.

***Figure S5. MET3 mutants do not show developmental defect***

(a) *met3* mutants transmission in self progeny. +/+ represent wild-type, +/- heterozygotes and -/- homozygotes plants for the mutations. (b) Percentage of seed abortion in Col-0 and *met3* mutants (*met3-3* and *met3-4*) siliques. (c) Seed size measurement for Col-0 and *met3* mutants dry seeds. No significant differences observed. (d) Pictures of Col-0 and *met3* mutants dry seeds. The scale bars represent 1 mm. (e) Pictures of Col-0 and *met3* mutants rosette at first (top panel) and fifth generation (bottom panel). The scale bars represent 1 cm. (f) Rosette area measurement in mm^2^ of Col and *met3* mutants at fifth generation. n=12. (g) Flowering time (number of days between transplanting and bolting) of Col and *met3* mutants at fifth generation. n=12.

***Figure S6. Transcriptome of met3 mutant seeds at 3DAP***

(a-b) Volcano plot depicting one differentially expressed gene comparing Col-0 to *met3-3* (a) and *met3-4* (*b*) at 3DAP.

