## Supplementary material for "DNA METHYLTRANSFERASE 3 (MET3) is regulated by Polycomb Group complex during Arabidopsis endosperm development": supfig

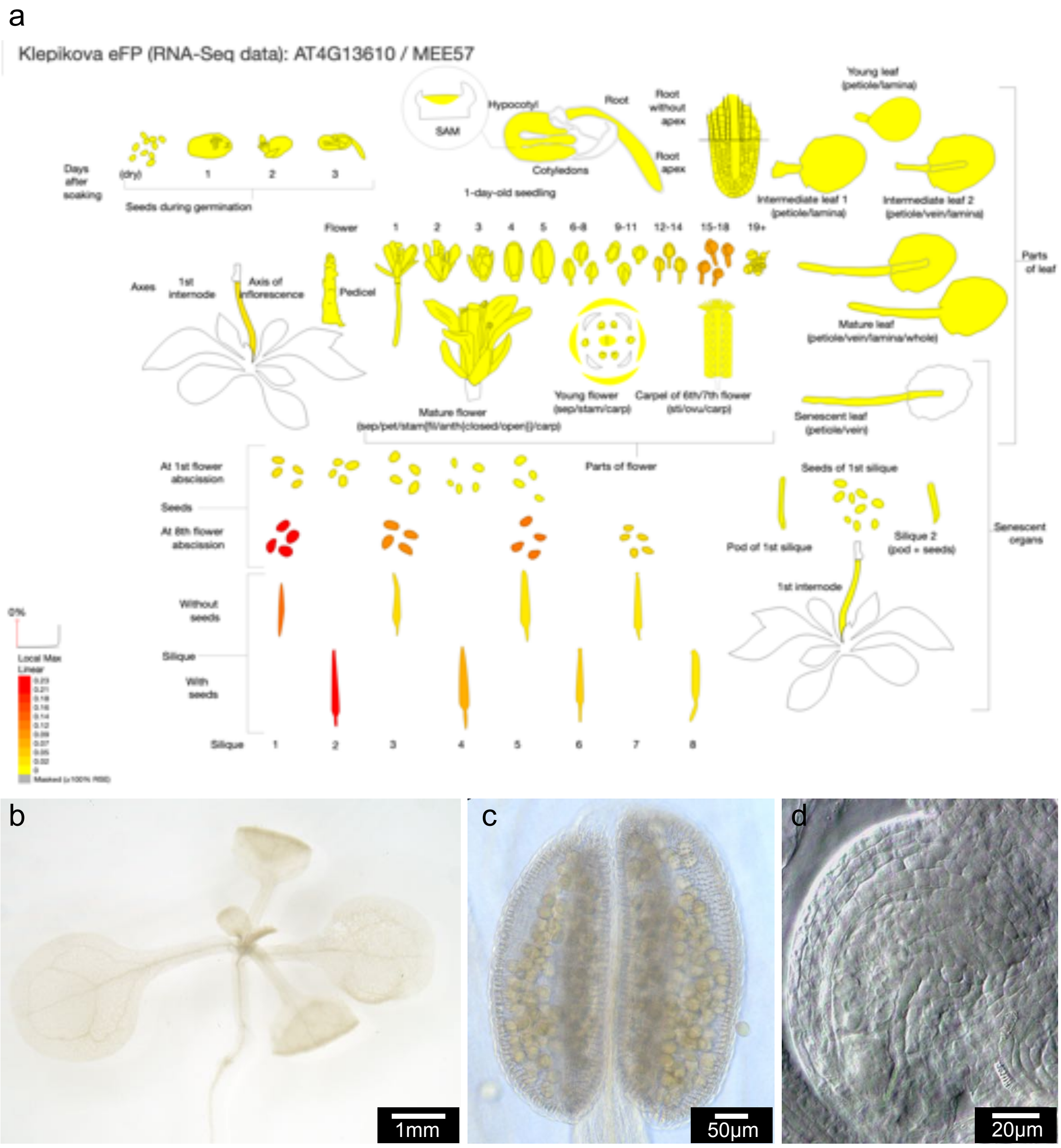


Figure S1. *MET3* is only expressed in seeds

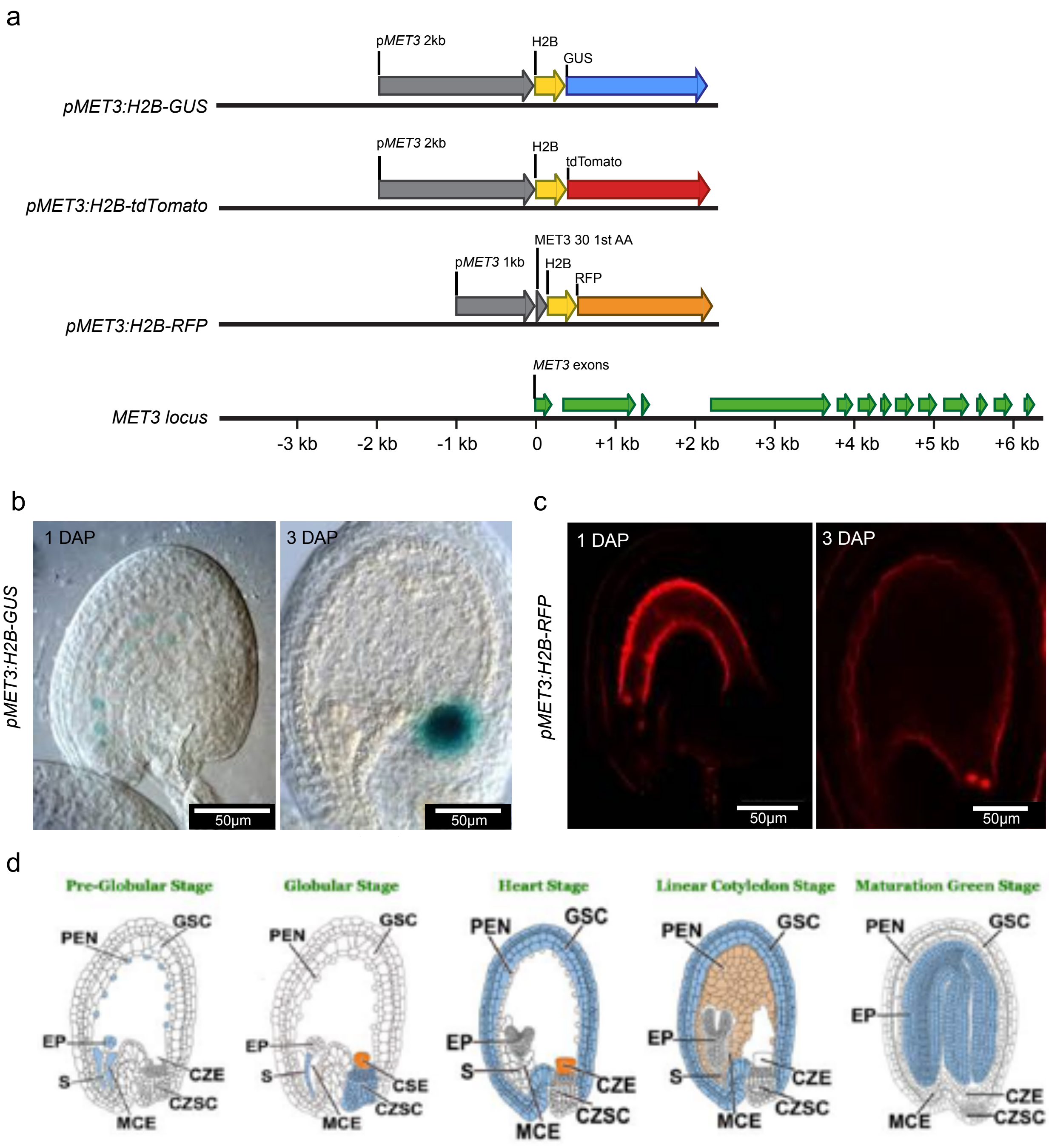


Figure S2. *MET3* reporters and expression

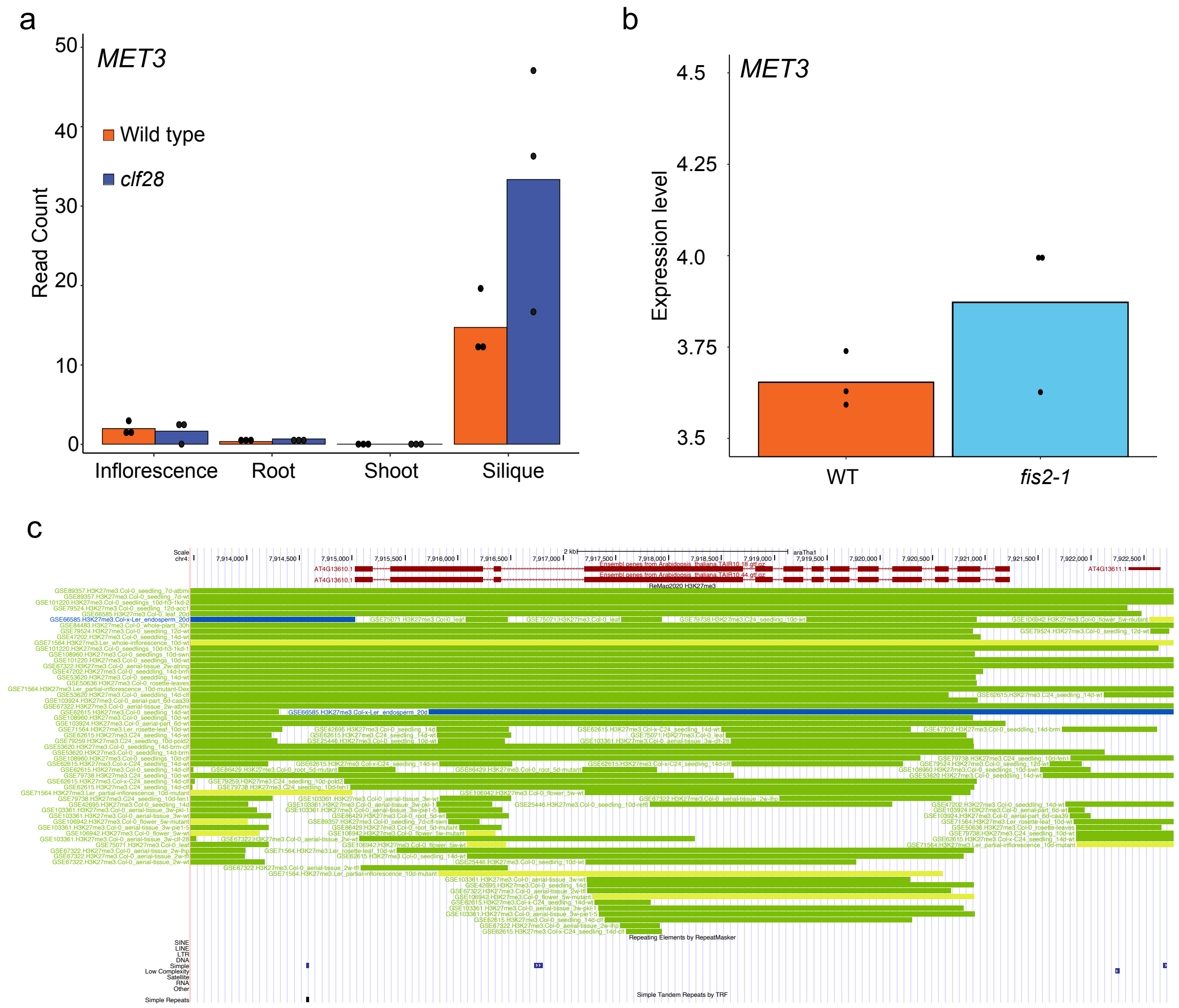


Figure S3. *MET3* regulation by Polycomb group

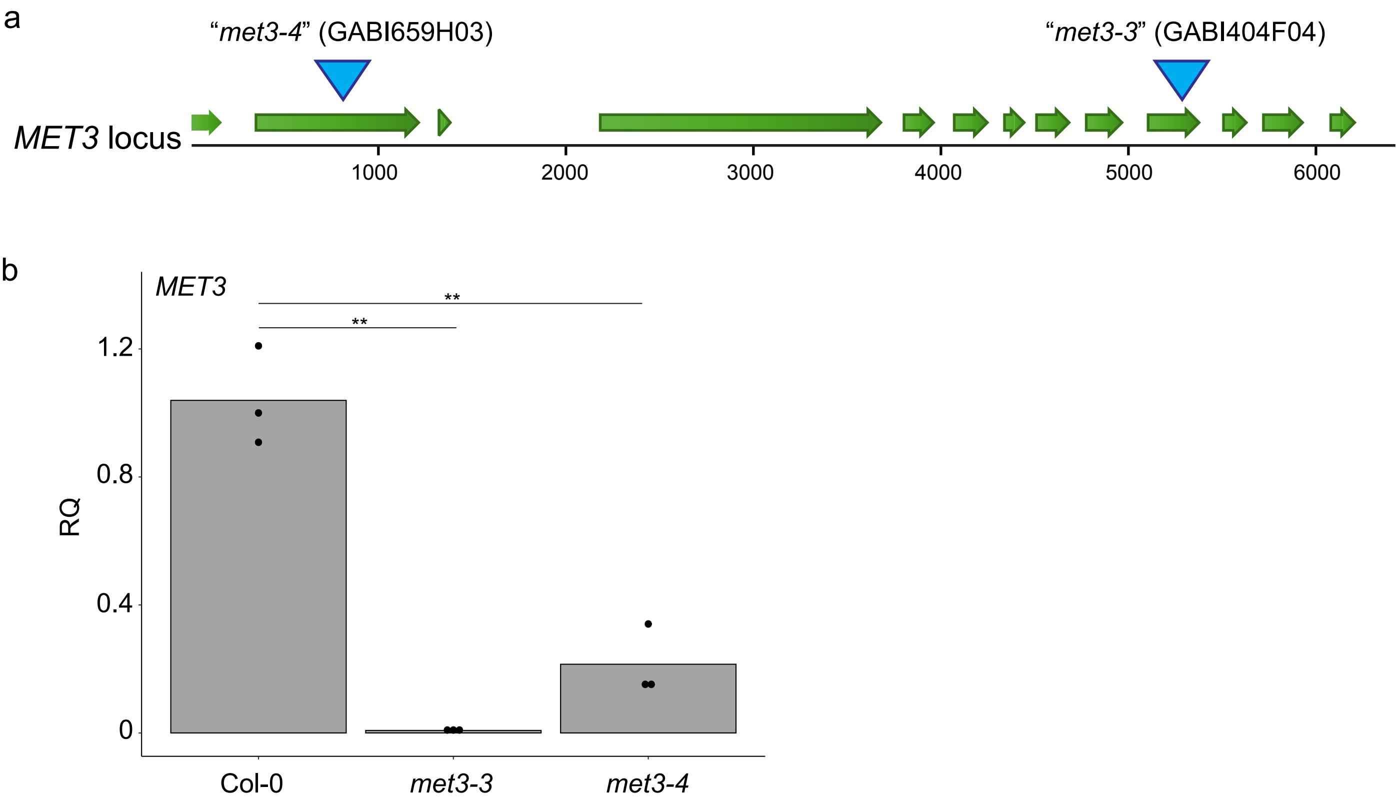


Figure S4. Characterization of *MET3* mutants

a

**Transmission rate**

| Progeny genotype | +/+ | -/+ | -/- | Total | Chi Square score |
|---|---|---|---|---|---|
| *met3-3* | 44 (25%) | 97 (55.11%) | 35 (19.88%) | 176 (100%) | 0.207 |
| *met3-4* | 39 (23.07%) | 85 (50.09%) | 45 (26.62%) | 169 (100%) | 0.806 |

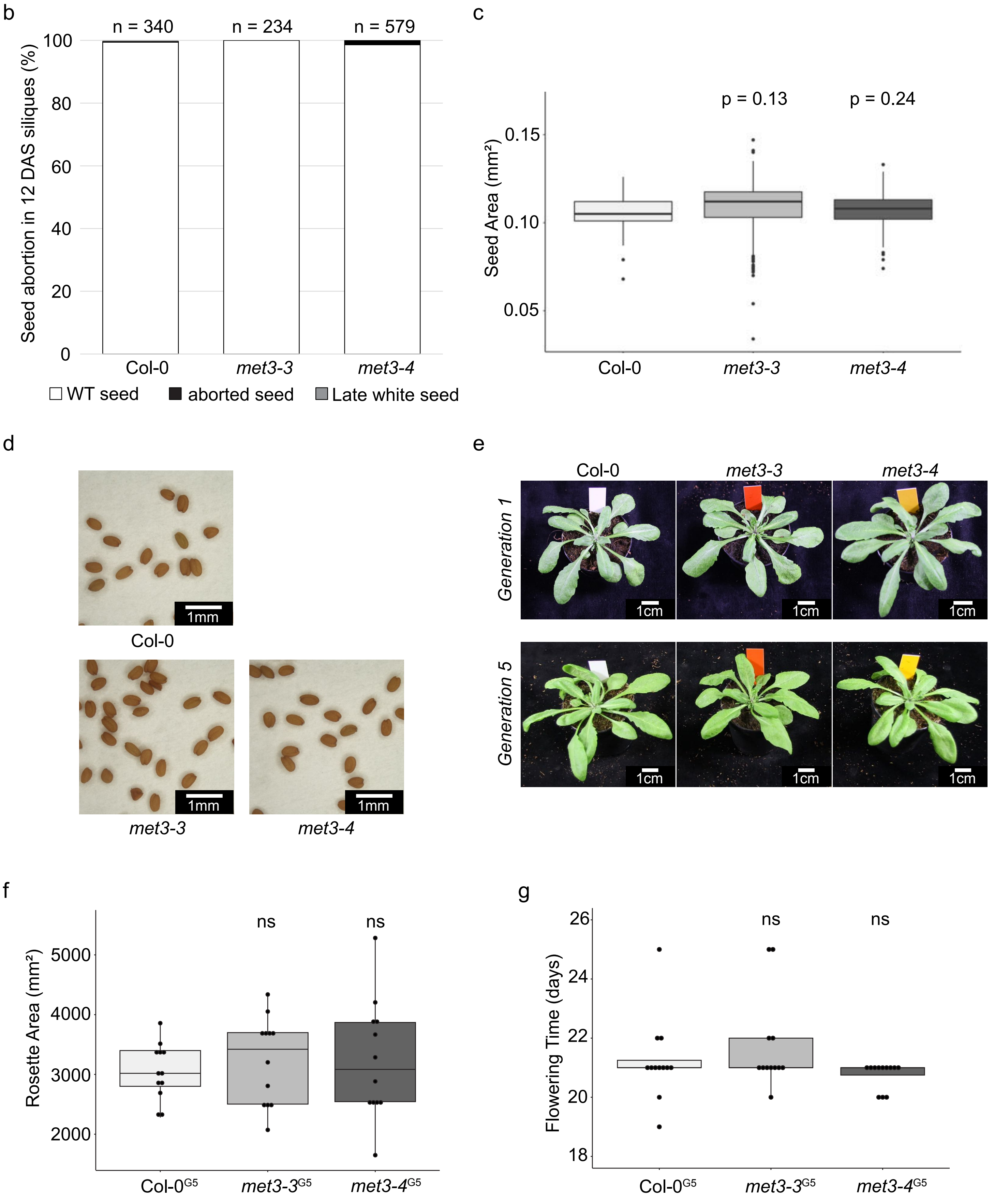

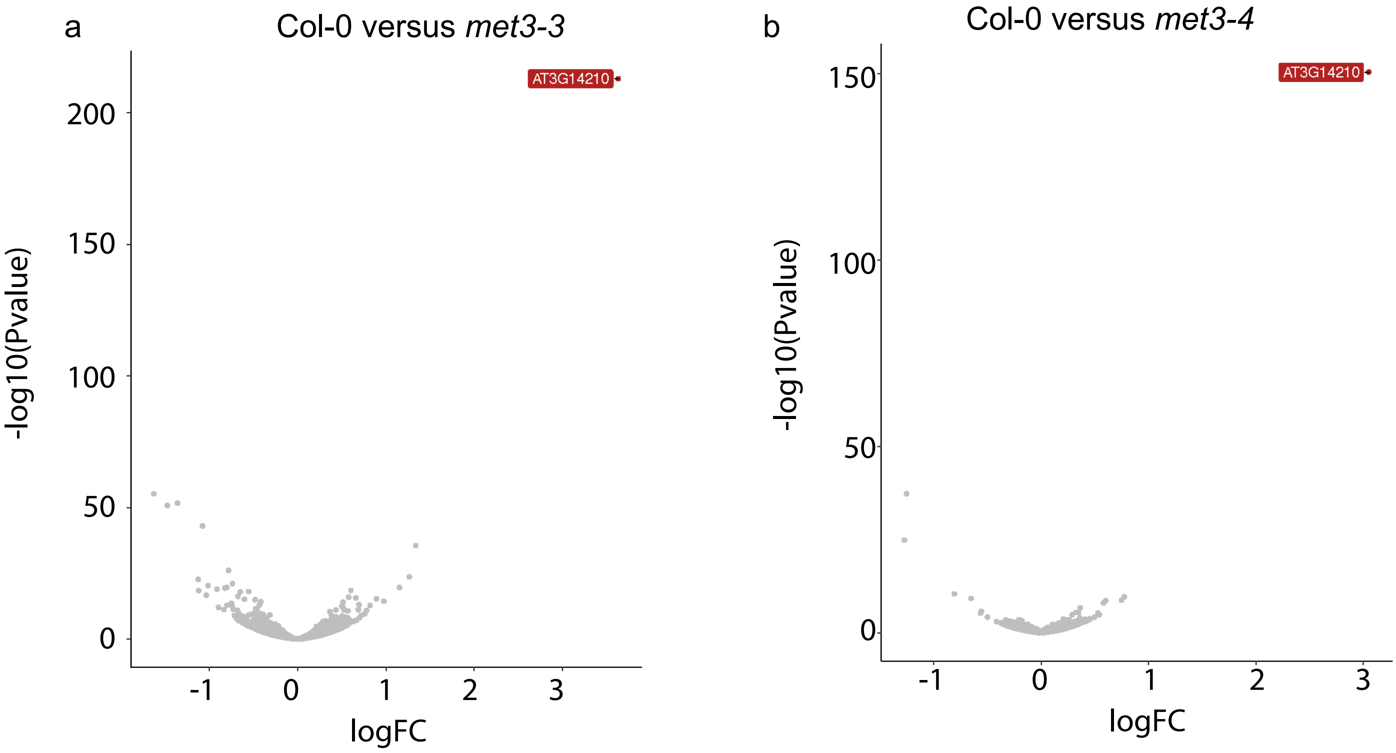


Figure S6. Transcriptome of *met3* mutant seeds at 3DAP
